# Similarity-based fusion of MEG and fMRI reveals spatio-temporal dynamics in human cortex during visual object recognition

**DOI:** 10.1101/032656

**Authors:** Radoslaw Martin Cichy, Dimitrios Pantazis, Aude Oliva

## Abstract

Every human cognitive function, such as visual object recognition, is realized in a complex spatio-temporal activity pattern in the brain. Current brain imaging techniques in isolation cannot resolve the brain’s spatio-temporal dynamics because they provide either high spatial or temporal resolution but not both. To overcome this limitation, we developed a new integration approach that uses representational similarities to combine measurements from different imaging modalities – magnetoencephalography (MEG) and functional MRI (fMRI) - to yield a spatially and temporally integrated characterization of neuronal activation. Applying this approach to two independent MEG-fMRI data sets, we observed that neural activity first emerged in the occipital pole at 50-80ms, before spreading rapidly and progressively in the anterior direction along the ventral and dorsal visual streams. These results provide a novel and comprehensive, spatio-temporally resolved view of the rapid neural dynamics during the first few hundred milliseconds of object vision. They further demonstrate the feasibility of spatially unbiased representational similarity based fusion of MEG and fMRI, promising new insights into how the brain computes complex cognitive functions.

## INTRODUCTION

A major challenge of cognitive neuroscience is to map the diverse spatio-temporal neural dynamics underlying cognitive functions under the methodological limitations posited by current neuroimaging technologies. Non-invasive neuroimaging technologies such as functional magnetic resonance imaging (fMRI) and magneto- and electroencephalography (M/EEG) offer either high spatial or high temporal resolution, but not both simultaneously. A comprehensive large-scale view of brain function with both high spatial and temporal resolution thus necessitates integration of available information from multiple brain imaging modalities, most commonly fMRI and M/EEG (for review see: (Dale and Halgren 2001; Debener et al. 2006; Rosa et al. 2010; Huster et al. 2012; Jorge et al. 2014)).

Here, we provide a novel approach to combine MEG and fMRI based on two basic principles. First, we assume *representational similarity:* if neural representations of two conditions are similarly represented in fMRI, they should also be similarly represented in MEG (Kriegeskorte 2008; Kriegeskorte and Kievit 2013; Cichy et al. 2014). Second, we assume *locality* of neural representation: information is represented in neuronal populations in locally restricted cortical regions, rather than being distributed across the whole brain. Based on these assumptions, linking the similarity relations in MEG for each millisecond with the similarity relations in a searchlight-based fMRI analysis (Haynes and Rees 2005; Kriegeskorte et al. 2006) promises a a spatio-temporally resolved account of neural activation.

We used the proposed methodology to yield a novel characterization of the spatiotemporal neural dynamics underlying a key cognitive function: visual object recognition. Visual object recognition recruits a temporally ordered cascade of neuronal processes (Robinson and Rugg 1988; Schmolesky et al. 1998; Bullier 2001; Cichy et al. 2014) in the ventral and dorsal visual streams (Ungerleider and Mishkin 1982; Milner and Goodale 2008; Kravitz et al. 2011). Both streams consist of multiple brain regions (Felleman and Van Essen 1991; Grill-Spector and Malach 2004; Wandell et al. 2007; Beeck et al. 2008; Grill-Spector and Weiner 2014) that encode different aspects of the visual input in neuronal population codes (Haxby et al. 2001; Pasupathy and Connor 2002; Haynes and Rees 2006; Kiani et al. 2007; Meyers et al. 2008; Kreiman 2011; Konkle and Oliva 2012; Tong and Pratte 2012; Cichy et al. 2013).

Analysis of two MEG-fMRI data sets from independent experiments yielded converging results: Visual representations emerged at the occipital pole of the brain early (~50-80ms), before spreading rapidly and progressively along the visual hierarchies in the dorsal and ventral visual stream, resulting in prolonged activation in high-level ventral visual areas. This result for the first time unravels the neural dynamics underlying visual object recognition in the human brain with millisecond and millimeter resolution in the dorsal stream, and corroborates previous findings regarding the successive engagement of the ventral visual stream in object representations, available until now only from intracranial recordings.

In sum, our results offer a novel description of the complex neuronal processes underlying visual recognition in the human brain, and demonstrate the efficacy and power and power of a similarity-based MEG-fMRI fusion approach to provide a refined spatio-temporally resolved view of brain function.

## MATERIALS AND METHODS

### Participants

We conducted two independent experiments. 16 right-handed, healthy volunteers (10 female, age: mean ± s.d. = 25.87 ± 5.38 years) participated in experiment 1, and 15 (5 female, age: mean ± s.d. = 26.60 ± 5.18 years) in experiment 2. All participants were right-handed with normal or corrected-to-normal vision and provided written consent. The study was conducted according to the Declaration of Helsinki and approved by the local ethics committee (Institutional Review Board of the Massachusetts Institute of Technology).

### Experimental design and task

Experimental design and task were similar for experiments 1 and 2. The stimulus set for experiment 1 was *C* = 92 real-world object images on a gray background, including human and animal faces, bodies, as well as natural and artificial objects (Figure 2A). The stimulus set for experiment 2 was *C* = 118 real-world object images on real backgrounds (Figure 4A). In both experiments, images were presented at the center of the screen at 2.9° (Experiment 1) and 4.0° (Experiment 2) visual angle with 500 ms duration, overlaid with a gray fixation cross. We adapted presentation parameters to the specific requirements of each acquisition technique.

For MEG, participants completed two sessions of 10 to 15 runs of 420 s each for experiment 1, and one session of 15 runs of 314 s each for experiment 2. Image presentation was randomized in each run. Trial onset asynchrony was 1.5 or 2 s (Experiment 1, Supplementary Figure 1) and 0.9-1 s (Experiment 2). Participants were instructed to respond to the image of a paper clip shown randomly every 3-5 trials (average 4) with an eye blink and a button press.

For fMRI, each participant completed two sessions. For experiment 1, each session consisted of 10-14 runs of 384 s each, and for experiment 2 each session consisted of 9-11 runs of 486 s each. Every image was presented once per run and image order was randomized with the restriction that the same condition was not presented on consecutive trials. We randomly interspersed 30 (experiment 1) or 39 (experiment 2) null trials presenting only a gray background. During null trials the fixation cross turned darker for 100ms and participants were instructed to respond to the change in luminance with a button press. TOS was 3 s, or 6 s with a preceding null trial.

### MEG acquisition

We acquired MEG signals from 306 channels (204 planar gradiometers, 102 magnetometers, Elekta Neuromag TRIUX, Elekta, Stockholm) at a sampling rate of 1 kHz, filtered between 0.03 and 330 Hz. We applied temporal source space separation (maxfilter software, Elekta, Stockholm (Taulu et al. 2004; Taulu and Simola 2006)) before analyzing data with Brainstorm (Tadel et al. 2011). For each trial we extracted peri-stimulus data from −100ms to +700ms, removed baseline mean and smoothed data with a 20-ms sliding window. We obtained 20-30 trials for each condition, session, and participant.

Next, we determined the similarity relations between visual representations as measured with MEG by multivariate pattern analysis (Figure 1). The rationale of using machine classifiers is that the better a classifier predicts condition labels based on patterns of MEG sensor measurements, the more dissimilar the MEG patterns are and thus the underlying visual representations. As such, classifier performance can be interpreted as a dissimilarity measure.

**Figure 1.**
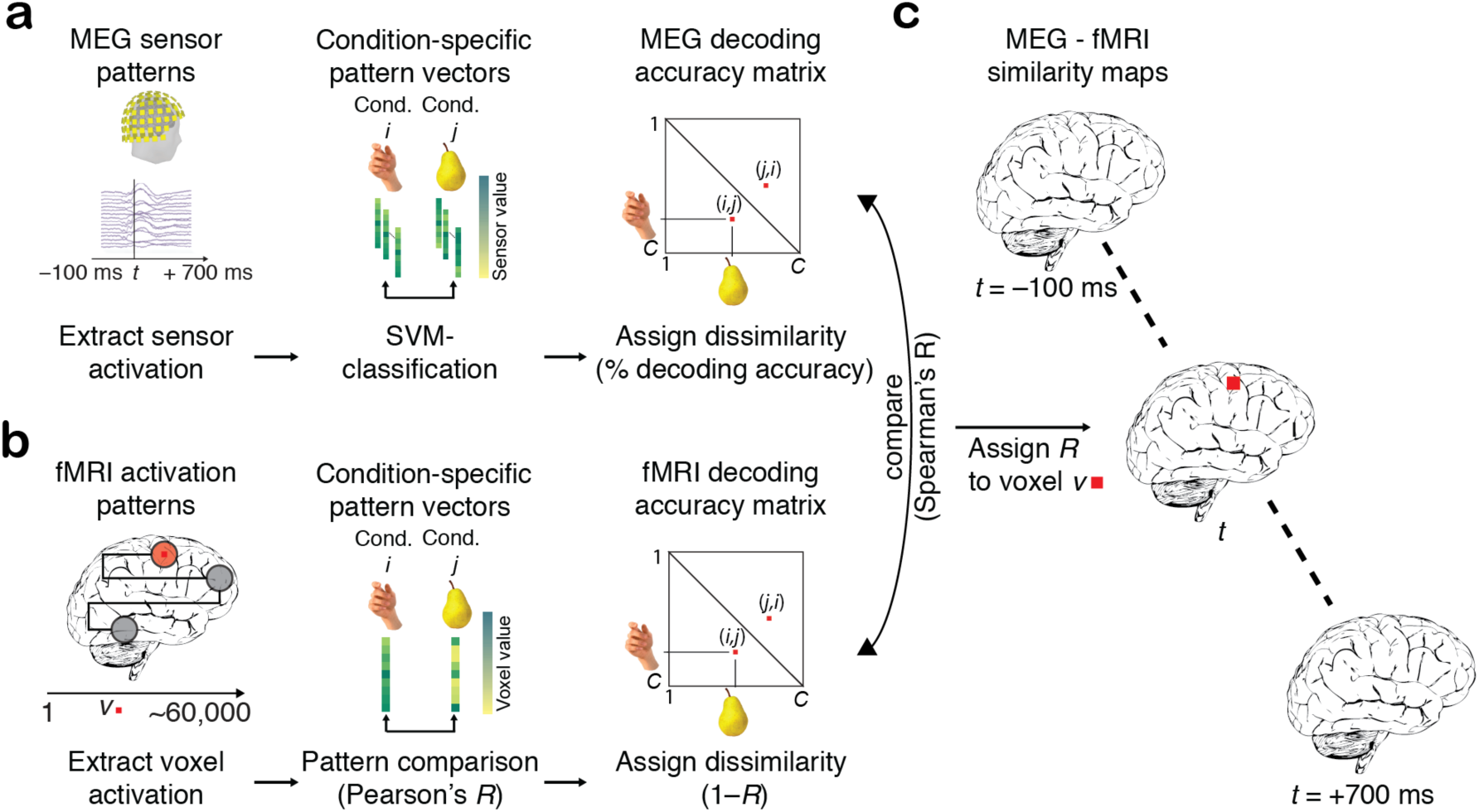
Spatially unbiased fMRI-MEG fusion analysis scheme, **a)** MEG analysis. We analyzed MEG data in a time-point specific fashion for each millisecond *t* from −100 to +700ms with respect to stimulus onset. For each pair of conditions, we used support vector machine classification to determine how well conditions were discriminated by multivariate MEG sensor patterns. Repeating this procedure for each condition pair yielded a condition-by-condition (*C* × *C*) matrix of decoding accuracies, constituting a summary of representational dissimilarities for a time point (MEG representational dissimilarity matrix, abbr. MEG RDM). **b)** fMRI analysis. We used searchlight analysis to reveal representational dissimilarity in locally constrained fMRI activity patterns. In detail, for each voxel *v* in the brain we extracted fMRI patterns in its local vicinity (4-voxel radius) and calculated condition-wise dissimilarity (1 – Spearman’s *R*), resulting in a condition-by-condition (*C × C*) fMRI representational dissimilarity matrix (fMRI RDM), constituting a summary of representational dissimilarities for a voxel’s vicinity. **c)** fMRI-MEG fusion. In the space of representational dissimilarities, MEG and fMRI results become directly comparable. For each time-specific MEG RDM, we calculated the similarity (Spearman’s R) to each fMRI searchlight’s fMRI RDM, yielding a 3D fMRI-MEG similarity map. Repeating this procedure for each millisecond yielded a movie of representational similarity correspondence in the brain, revealing spatiotemporal neural dynamics.

We conducted the multivariate analysis in the form of linear support vector machine (SVM) classification independently for each subject and session. For each peristimulus time point from −100 to +700ms, preprocessed MEG was extracted and arranged as 306 dimensional measurement vectors (corresponding to the 306 MEG sensors), yielding *M* pattern vectors per time point and condition (image). We then used supervised learning with a leave-one-trial out cross validation to train a SVM classifier (LibSVM implementation, www.csie.ntu.edu.tw/~cjlin/libsvm) to pairwise discriminate any two conditions. For each time point and condition pair, *M*-1 measurement vectors were assigned to the training set and used to train a support vector machine. The left out *M*th vectors for the two conditions were assigned to a testing set and used to assess the classification performance of the trained classifier (percent decoding accuracy). The training and testing process was repeated 100 times with random assignment of trials to the training and testing set. Decoding results were averaged across iterations, and the average decoding accuracy was assigned to a matrix of size *C × C* (*C* = 92 for experiment 1, 118 for experiment 2), with rows and columns indexed by the classified conditions. The matrix was symmetric and the diagonal was undefined. This procedure yielded one *C* × *C* matrix of decoding accuracies for every time point, referred to as MEG representational dissimilarity matrix (MEG RDM). For experiment 2 measurement vectors were averaged by 5 before entering multivariate analysis to reduce computational load.

### fMRI acquisition

MRI scans were conducted on a 3T Trio scanner (Siemens, Erlangen, Germany) with a 32-channel head coil. In both experiments and each session, we acquired structural images using a standard T1-weighted sequence (192 sagittal slices, FOV = 256 mm^2^, TR = 1,900 ms, TE = 2.52 ms, flip angle = 9°).

Functional data were collected with two different protocols. In experiment 1 (partial brain coverage) data had high spatial resolution, but covered the ventral visual brain only. In each of two sessions we acquired 10–14 runs of 192 volumes each for each participant (gradient-echo EPI sequence: TR = 2,000 ms, TE = 31 ms, flip angle = 80°, FOV read = 192 mm, FOV phase = 100%, ascending acquisition, gap = 10%, resolution = 2 mm isotropic, slices = 25). The acquisition volume covered the occipital and temporal lobe and was oriented parallel to the temporal cortex.

The second data set (experiment 2) had lower spatial resolution, but covered the whole brain. We acquired 9-11 runs of 648 volumes for each participant (gradientecho EPI sequence: TR = 750 ms, TE = 30 ms, flip angle = 61°, FOV read = 192 mm, FOV phase = 100% with a partial fraction of 6/8, through-plane acceleration factor 3, bandwidth 1816Hz/Px, resolution = 3mm^3^, slice gap 20%, slices = 33, ascending acquisition).

### fMRI analysis

We preprocessed fMRI data using SPM8 (http://www.fil.ion.ucl.ac.uk/spm/). Analysis was identical for the two experiments. For each participant, fMRI data were realigned and co-registered to the T1 structural scan acquired in the first MRI session. Then, MRI data was normalized to the standard MNI template. We used a general linear model (GLM) to estimate the fMRI response to the 92 (experiment 1) or 118 (experiment 2) image conditions. Image onsets and duration entered the GLM as regressors and were convolved with a hemodynamic response function. Movement parameters were included as nuisance parameters. Additional regressors modeling the two sessions were included in the GLM. The estimated condition-specific GLM parameters were converted into *t*-values by contrasting each condition estimate against the implicitly modeled baseline.

We then analyzed fMRI data in a spatially unbiased approach using a searchlight analysis method (Kriegeskorte et al. 2006; Haynes et al. 2007). We processed each subject separately. For each voxel v, we extracted condition-specific *t*-value patterns in a sphere centered at v with a radius of 4 voxels (searchlight at *v*) and arranged them into fMRI *t*-value pattern vectors. For each pair of conditions we calculated the pair-wise dissimilarity between pattern vectors by 1 minus Spearman’s R, resulting in a 92 × 92 (experiment 1) or 118 × 118 (experiment 2) fMRI representational dissimilarity matrix (fMRI RDM) indexed in columns and rows by the compared conditions. fMRI RDMs were symmetric across the diagonal, and entries were bounded between 0 (no dissimilarity) and 2 (complete dissimilarity). This procedure resulted in one fMRI RDM for each voxel in the brain.

### MEG - fMRI fusion with representational similarity analysis

To relate neuronal temporal dynamics observed in MEG with their spatial origin as indicated by fMRI, we used representational similarity analysis (Kriegeskorte 2008; Kriegeskorte and Kievit 2013; Cichy et al. 2014). The basic idea is that if two images are similarly represented in MEG patterns, they should also be similarly represented in fMRI patterns. Comparing similarity relations in this way allows linking particular locations in the brain to particular time points, yielding a spatio-temporally resolved view of the emergence of visual representations in the brain.

We compared fMRI RDMs with MEG RDMs. For experiment 1, analysis was conducted within subject, comparing subject-specific fMRI and MEG RDMs. For experiment 2, only one MEG session was conduced, and to increase power we first averaged MEG RDMs across participants before comparing the subject-averaged MEG RDMs with the subject-specific fMRI RDMs.

Further analysis was identical for both experiments and independent for each subject. For each time point we computed the similarity (Spearman’s R) between the MEG RDM and the fMRI RDM of each voxel. This yielded a 3D map of representational similarity correlations, indicating locations in the brain at which neuronal processing emerged at a particular time point. Repeating this analysis for each time point yielded a spatio-temporally resolved view of neural activity in the brain during object perception.

### Statistical testing

We used permutation tests for random-effects inference and corrected results for multiple comparisons with a cluster level correction (Nichols and Holmes 2002; Pantazis et al. 2005; Maris and Oostenveld 2007). To determine a cluster-definition threshold, we aggregated subject-averaged data across voxels and time points from −100 to 0ms and determined the 99.9% confidence intervals. This constitutes a baseline-based cluster definition threshold at *p* = 0.001.

We then randomly shuffled the sign of subject-specific data points (1000 times), averaged data across subjects, and determined 4-dimensional clusters by spatial and temporal contiguity at the cluster definition threshold. Storing the maximal cluster size for each resample yielded a distribution of cluster sizes under the null hypothesis. We report clusters as significant if they were greater than the 95% threshold defined on the null cluster size distribution (i.e. cluster-definition threshold *p* =0.05).

## RESULTS

### MEG-fMRI fusion in representational similarity space

In two independent experiments, participants viewed images of real world objects: 92 objects on a gray background in experiment 1 (Figure 2a), and 118 objects on a real-world background in experiment 2 (Figure 4a). Each experiment consisted of separate recording sessions presenting the same stimulus material during MEG or fMRI data collection. Images were presented in random order in experimental designs optimized for each imaging modality. Subjects performed orthogonal vigilance tasks.

**Figure 2.**
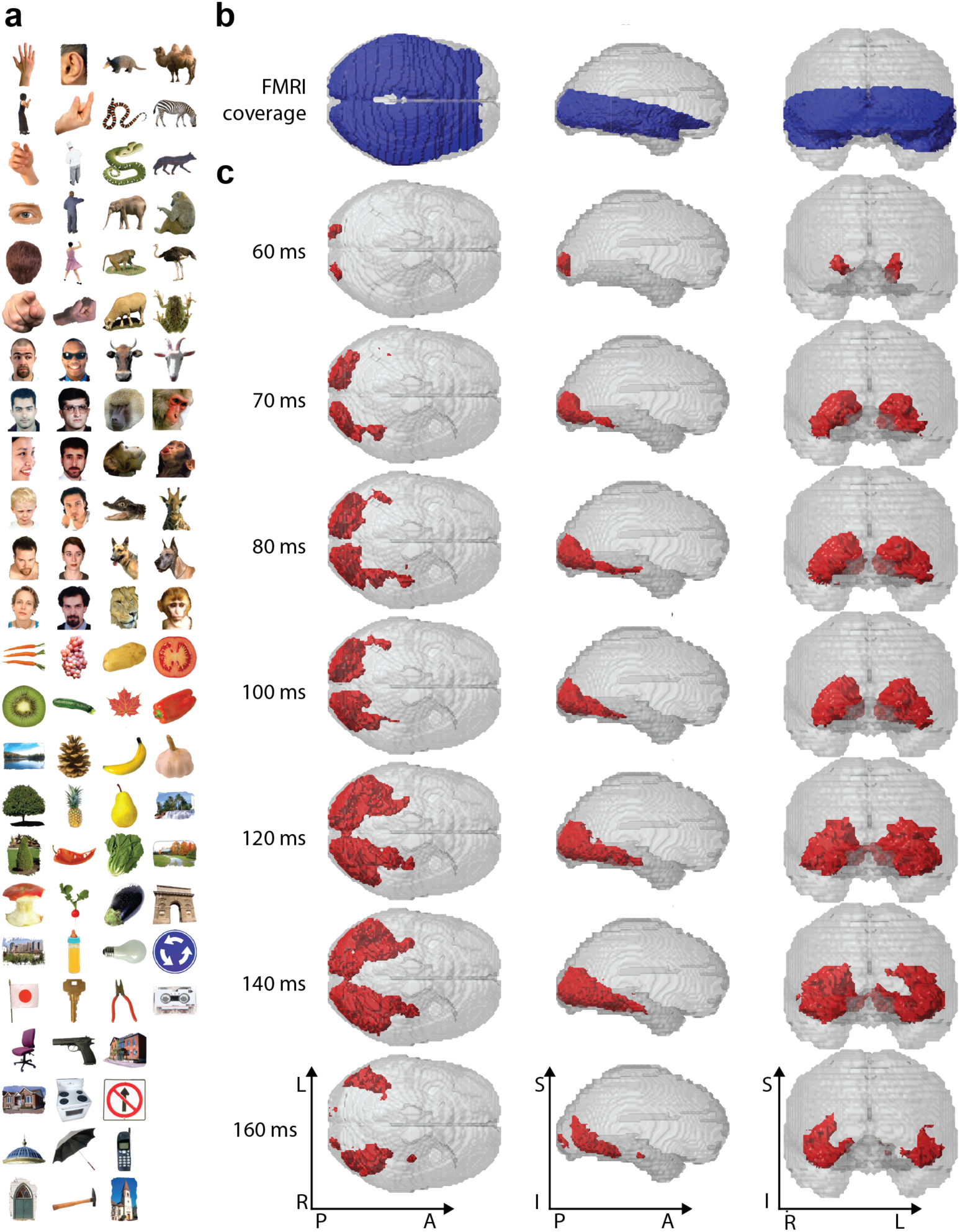
Stimulus material, fMRI brain coverage and significant MEG-fMRI representational similarity correspondences for experiment 1. **a)** The stimulus set consisted of 92 cropped objects, including human and animal bodies and faces, as well as natural and artificial objects. **b)** We recorded fMRI with partial brain coverage (blue regions), reflecting the trade-off between high-resolution (2 mm isovoxel), extent of coverage, and temporal sampling (TR=2s). **c)** MEG-fMRI fusion revealed spatio-temporal neural dynamics in the ventral visual stream, originating at the occipital lobe and extending rapidly in anterior direction along the ventral visual stream. Red voxels indicate statistical significance (*n*=15, cluster-definition threshold *p*<0.001, cluster threshold *p*<0.05). A millisecond resolved movie is available as Supplementary Movie 1.

To integrate MEG with fMRI data, we abstracted from the sensor space of each imaging modality to a common similarity space, defined by the similarity of condition-specific response patterns in each modality. Our approach relies on representational similarity analysis (Kriegeskorte 2008; Cichy et al. 2014), and effectively constructs MEG and fMRI representational similarity relations that are directly comparable.

In detail, for MEG we analyzed data in a time-point specific fashion for each millisecond from −100 to +1,000ms with respect to stimulus onset (Figure 1). To make full use of the information encoded at each time point across MEG sensors, we used multivariate pattern analysis (Carlson et al. 2013; Cichy et al. 2014; Clarke et al. 2014; Isik et al. 2014). Specifically, for each time point we arranged the measurements of the entire MEG sensor array into vectors, resulting in different patterns vectors for each condition, stimulus repetition, and time point. These pattern vectors were the input to a support vector machine procedure designed to classify different pairs of conditions (images). Repeating the classification procedure for all pairs of conditions yielded a condition × condition (*C* × *C*) matrix of classification accuracies, one per time point. Interpreting classification accuracies as a representational dissimilarity measure, the *C* × *C* matrix summarizes for each time point which conditions are represented similarly (low decoding accuracy) or dissimilarly (high decoding accuracy). The matrix is thus termed MEG representational dissimilarity matrix (MEG RDM).

For fMRI, we used a searchlight approach (Haynes and Rees 2005; Kriegeskorte et al. 2006) in combination with multivariate pattern similarity comparison (Kriegeskorte 2008; Kriegeskorte et al. 2008; Kriegeskorte and Kievit 2013) to extract information stored in local neural populations resolved with the high spatial resolution of fMRI. For every voxel in the brain, we extracted condition-specific voxel patterns in its vicinity, and calculated pair-wise dissimilarity (1 minus Pearson’s R) between voxel patterns of different conditions. Repeating this procedure for all pairs of conditions yielded a *C* × *C* fMRI representational dissimilarity matrix (fMRI RDM) summarizing for each voxel in the brain which conditions are represented similarly or dissimilarly.

Thus, MEG and fMRI data became directly comparable to each other in similarity space via their RDMs, allowing integration of their respective high temporal and spatial resolution. For a given time point, we compared (Spearman’s *R*) the corresponding MEG RDM with each searchlight-specific fMRI RDM, storing results in a 3-dimensional (3D) volume at the location of the center voxel of each searchlight. Each time-specific 3D volume indicated where in the brain the fMRI representations were similar to the ongoing MEG representations. Repeating this procedure for all time points yielded a set of brain volumes that unraveled the spatio-temporal neural dynamics underlying object recognition. We corrected for multiple comparisons across space and time by 4-dimensional cluster correction (cluster-definition threshold *p*<0.001, cluster threshold *p*<0.05).

### MEG-fMRI fusion reveals successive cortical region activation in the ventral and dorsal visual stream during visual object recognition

We applied the MEG-fMRI fusion method to data from experiment 1, i.e. visual responses to a set of *C* = 92 images of real world objects (Figure 2a). fMRI data had only a partial brain coverage of occipital and temporal cortex (Figure 2b). Results revealed the spatio-temporal dynamics in the ventral visual pathway in the first few hundred milliseconds of visual object perception. We observed the emergence of neuronal activity with onset around ~50-60ms at the occipital pole, succeeded by rapid activation in the anterior direction into the temporal lobe (Figure 2c, for ms-resolved see Supplementary Movie 1). A projection of the results onto axial slices at 140ms after stimulus onset exemplifies the results (Figure 3, for ms-resolved see Supplementary Movie 2).

**Figure 3.**
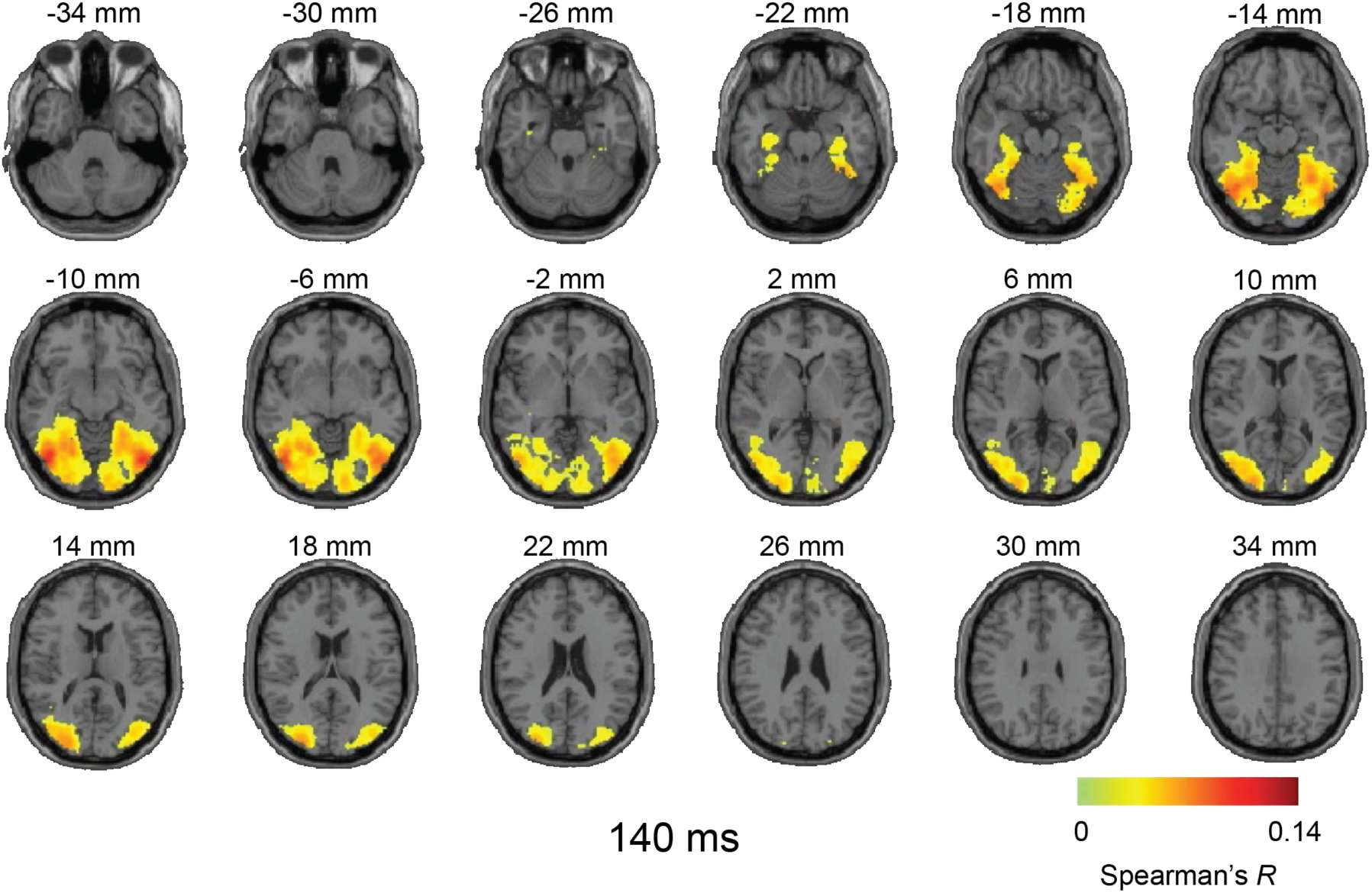
Axial slices exemplifying the MEG-fMRI fusion results for experiment 1 at 140ms. MEG-fMRI representation correlations were projected onto axial slices of a standard T1 image in MNI space. The results indicate neural activity reaching far into the ventral visual stream in temporal cortex. Color-coded voxels indicate strength of MEG-fMRI representation correlations (Spearman’s *R*, scaled between 0 and maximal observed value, *n*=15, cluster-definition threshold *p*<0.001, cluster threshold *p*<0.05). A millisecond resolved movie is available as Supplementary Movie 2.

Due to the limited MRI coverage in experiment 1, spatio-temporal dynamics beyond the ventral visual cortex, and in particular the dorsal visual stream, could not be assessed (Andersen et al. 1987; Sereno and Maunsell 1998; Sereno et al. 2002; Denys et al. 2004; Lehky and Sereno 2007; Janssen et al. 2008; Konen and Kastner 2008). To provide a full-brain view of spatio-temporal neuronal dynamics during object recognition, and to assess the reproducibility of the fMRI-MEG fusion method, we conducted experiment 2 with full brain MRI coverage (Figure 4b) and a different, more extended, set of object images (*C* = 118; Figure 4a). We again observed early neuronal activity in the occipital pole, though with a somewhat slower significant onset (~70-80ms), and rapid successive spread in the anterior direction across the ventral visual stream into the temporal lobe (Figure 4c, Supplementary Movie 3), reproducing the findings of experiment 1. Crucially, we also observed a progression of neuronal activity along the dorsal visual pathway into the inferior parietal cortex from about 140ms onward. Figure 5 shows example results at 180ms projected onto axial slices (for ms-resolved see Supplementary Movie 4).

**Figure 4.**
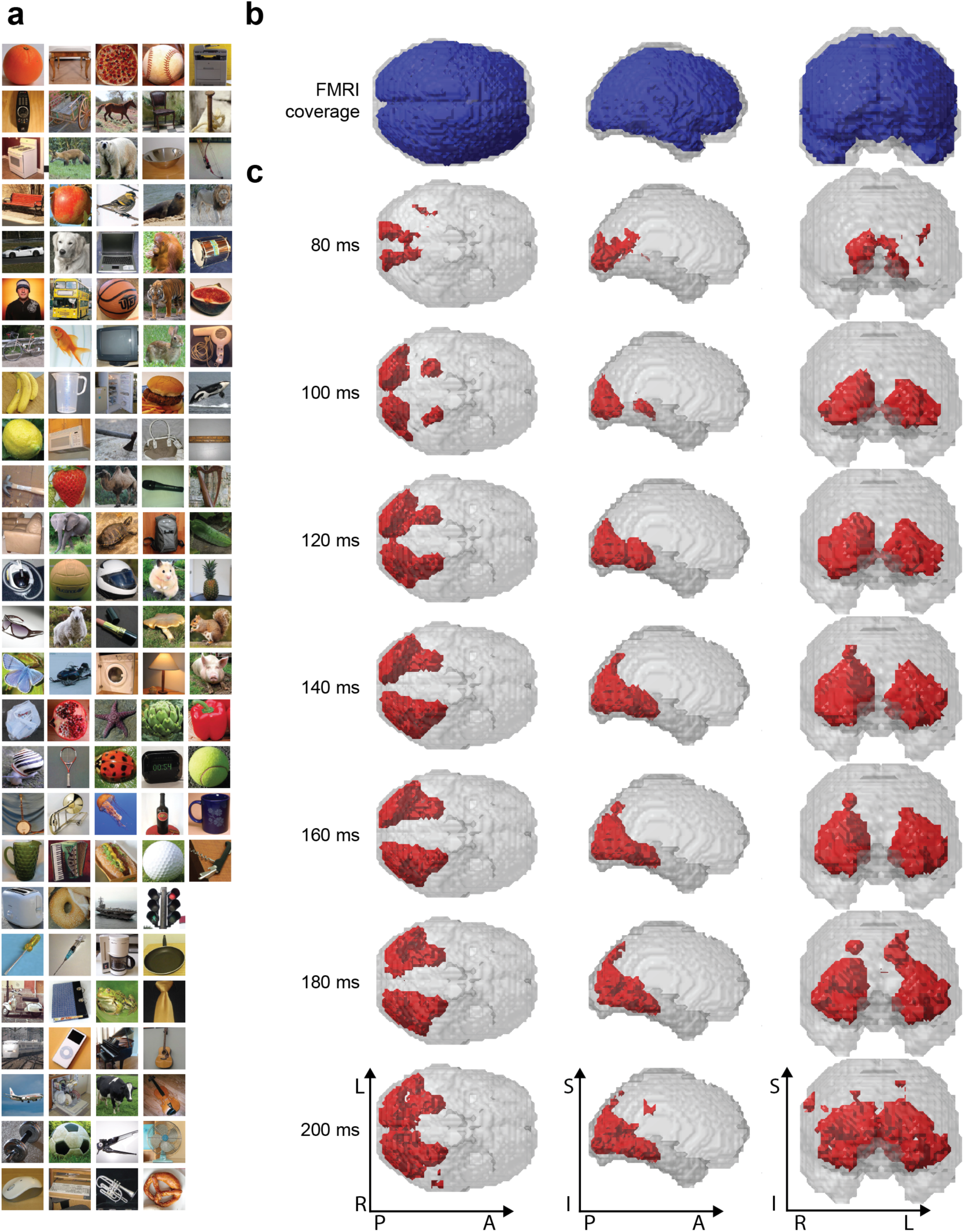
Stimulus material, brain coverage and significant MEG-fMRI representational similarity correspondences for experiment 2. **a)** The stimulus set consisted of 118 objects on natural background. **b)** We recorded fMRI with a whole-brain coverage (blue regions). **c)** Whole-brain analysis revealed neural dynamics along the dorsal visual stream into parietal cortex in addition to the ventral visual stream. Red voxels indicate statistical significance (*n*=15, clusterdefinition threshold *p*<0.001, cluster threshold *p*<0.05). A millisecond resolved movie is available as Supplementary Movie 3.

**Figure 5.**
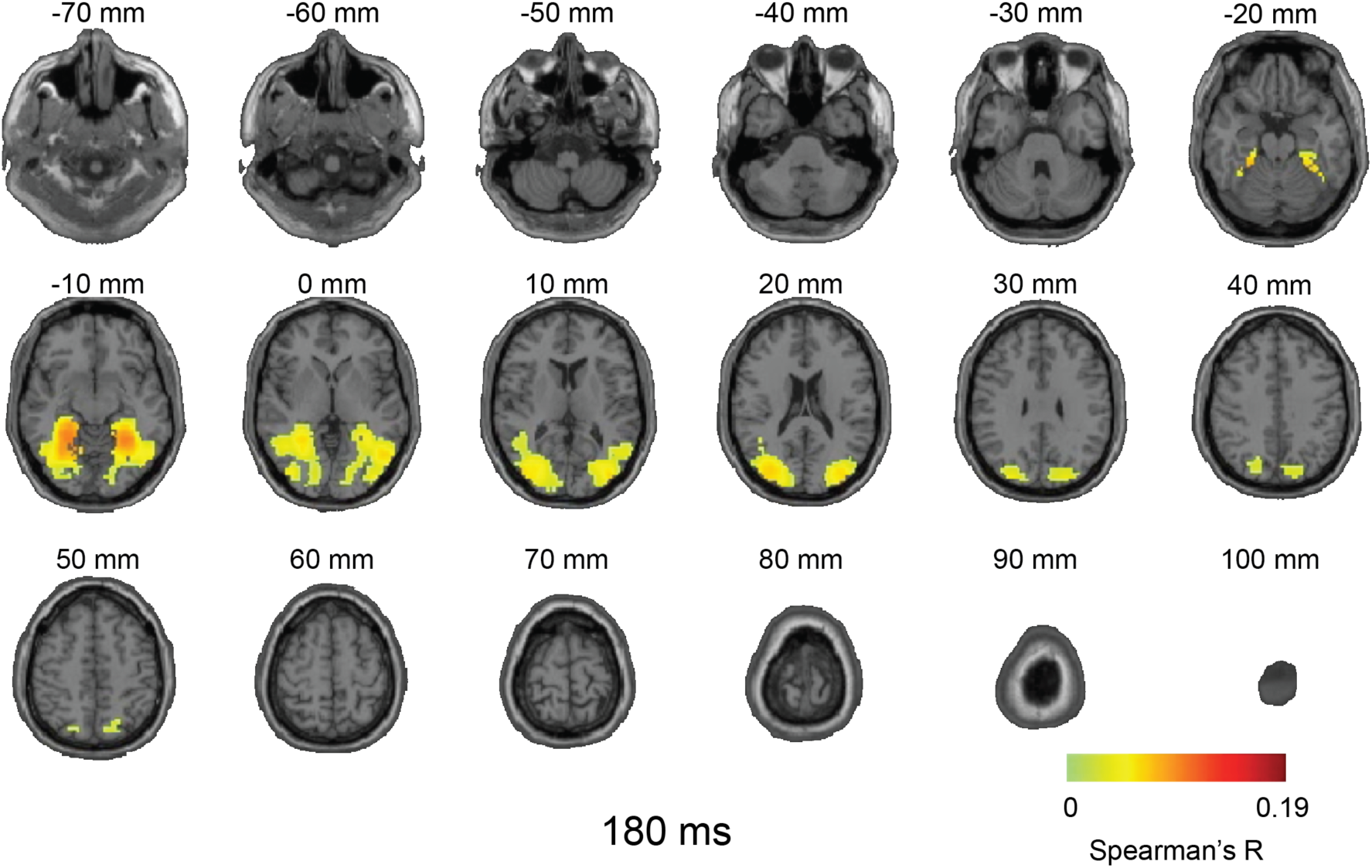
Axial slices exemplifying the MEG-fMRI fusion results for experiment 2 at 180ms. MEG-fMRI representation correlations were projected onto axial slices of a standard T1 image in MNI space. In addition to ventral cortex activation, reproducing experiment 1, we also observed activation far up the dorsal visual stream in inferior parietal cortex. Color-coded voxels indicate strength of MEG-fMRI representation correlations (Spearman’s *R*, scaled between 0 and maximal value observed, *n*=15, cluster-definition threshold *p*<0.001, cluster threshold *p*<0.05). A millisecond resolved movie is available as Supplementary Movie 4.

Together, our results describe the spatio-temporal dynamics of neural activation underlying visual object recognition during the first few hundred milliseconds of vision as a cascade of cortical region activations in both the ventral and dorsal visual pathways.

## DISCUSSION

### Summary

To resolve spatio-temporal neural dynamics in the human brain during visual object recognition, we proposed a MEG-fMRI fusion method based on representational similarities and multivariate pattern analysis. The locus of activation onset was at the occipital pole, followed by rapid and progressive activation along the processing hierarchies of the dorsal and ventral visual streams into temporal and parietal cortex respectively. This result provides a comprehensive and refined view of the complex spatio-temporal neural dynamics underlying visual object recognition in the human brain. It further suggests that the proposed methodology is a strong analytical tool for the study of any complex cognitive function.

### Tracking neural activity in visual object recognition

Our results revealed the spatio-temporal neural dynamics during visual object recognition in the human brain as neuronal activation emerged in the occipital lobe and rapidly spread along the ventral and dorsal visual pathways.

Concerning the dorsal visual stream, our results provide novel insights into the timing of neural activity in parietal cortex. To our knowledge, no gold-standard intracranial response profiles for parietal cortex have been determined in humans due to typically scarce electrode coverage and low visual responsiveness in that part of the cortex (Liu et al. 2009). In monkey, two measures of neuronal timing in parietal cortex have been assessed: onset and peak response latencies. While onset latencies were short and varied between 50-90ms (Robinson and Rugg 1988; Colby et al. 1996; Lehky and Sereno 2007; Janssen et al. 2008; Romero et al. 2012), peak latencies occurred later at 100-200ms, often followed by a long plateau of high firing rates (Colby et al. 1996; Shikata et al. 1996; Lehky and Sereno 2007; Janssen et al. 2008; Romero et al. 2012). The similarity of peak latency in electrophysiological recordings and the results observed here suggest that our approach reveals spatiotemporal neural dynamics mostly matching peak rather than onset firing rates in the dorsal stream.

Concerning the ventral stream, our results concur with previously reported increases of response latencies along the posterior-anterior gradient in the ventral pathway in humans (Mormann et al. 2008; Liu et al. 2009) and monkeys (Schmolesky et al. 1998; Bullier 2001). Together, our results provide evidence that object recognition is a hierarchical process unfolding over time (Robinson and Rugg 1988; Schmolesky et al. 1998; Bullier 2001; Cichy et al. 2014) and space (Ungerleider and Mishkin 1982; Felleman and Van Essen 1991; Grill-Spector and Malach 2004; Wandell et al. 2007; Beeck et al. 2008; Kravitz et al. 2011; Grill-Spector and Weiner 2014) in both dorsal and ventral streams (Ungerleider and Mishkin 1982; Milner and Goodale 2008; Kravitz et al. 2011).

### Relation of the approach to the standard taxonomy of fMRI-M/EEG integration approaches

How does our approach fit into the taxonomy of previous approaches for integrating fMRI and M/EEG, and in effect how does it relate to its advantages and limitations? Typically 4 distinct categories are proposed: asymmetric approaches where one modality constraints the other, i.e. (1) fMRI-constrained M/EEG and (2) M/EEG constrained fMRI analysis, and symmetric approaches (fusion) in which data from either modality is weighted equally, combined in a (3) model- or (4) data-driven way (for review see e.g. (Rosa et al. 2010; Huster et al. 2012; Sui et al. 2012)).

Our approach does not fit the asymmetric constraint taxa. fMRI does not constrain the solution of M/EEG source reconstruction algorithms, nor do M/EEG features enter into the estimation of voxel-wise activity. Instead, fMRI and MEG constrain each other symmetrically via representational similarity. Thus our approach evades the danger of asymmetric approaches in giving an unwarranted bias to either imaging modality.

By exclusion this suggests that our approach belongs to a symmetric approach taxon. However, our approach also differs critically from previous symmetric approaches, both model- and data-based. Whereas model-based approaches typically depend on an explicit generative forward model from neural activity to both fMRI and M/EEG measurements, our approach does not. The advantage of this is that our approach evades the computational complexity of solutions to such models, as well as the underlying necessary assumptions about the complex relationship between neuronal and BOLD activity (Logothetis and Wandell 2004; Logothetis 2008).

Data-driven symmetric approaches, in turn, typically utilize unsupervised methods such as independent component analysis for integration, whereas our approach relies on the explicit assumption of similarity to constrain the solution space. Our approach is data-driven because it uses correlations to extract and compare representational patterns. However it has the advantage that, whereas understanding results of most data-driven approaches need additional interpretational steps, our results are directly interpretable.

In summary, our approach is fundamentally different than existing fMRI and M/EEG integration approaches and offers the distinct advantages: while providing interpretational ease due to the explicit constraint of representational similarity, it evades the bias of asymmetric approaches as well as the assumptions and complexities of symmetric approaches.

### Sensitivity, specificity, and potential ambiguity of the proposed approach

Our approach has several characteristics that provide high sensitivity and specificity with a limited risk of ambiguity in the results. The first advantage is its high sensitivity to detect information in either modality through the use of multivariate analyses. Multivariate analyses of both fMRI activity patterns (Haxby et al. 2001; Haynes and Rees 2006; Kriegeskorte et al. 2006; Norman et al. 2006; Tong and Pratte 2012) and MEG sensor patterns (Carlson et al. 2013; Cichy et al. 2014; Clarke et al. 2014; Isik et al. 2014) have been shown to provide increased informational sensitivity over mass-univariate approaches.

While our integration approach epitomizes on the increased sensitivity provided by multivariate analysis in each modality, the multivariate treatment is not complete. Our approach does not allow estimation of the relation between any set of voxels and MEG data, but is constrained in that only local response fMRI patterns are considered in the searchlight approach. We thus trade the ability to account for widely distributed information across the whole brain for precise localization and computational tractability.

The second advantage of the integration approach is the high specificity of the integration results. By seeking shared MEG and fMRI similarity patterns across a large set of conditions, i.e. covariance in similarity relations, our approach provides rich constraints on integration: for a set of any *C* conditions, there are ((*C* × *C*)−*C*)/2 different values in the representation matrices being compared. In contrast, previous approaches relied on smaller sets, i.e. co-variance in activation across few steps of parametric modulation (Mangun et al. 1997; Horovitz et al. 2004; Schicke et al. 2006), or co-variance across subjects (Sadeh et al. 2011).

However, the high specificity of our approach is limited to experimental contexts where a large number of conditions are available. For best results, specifically optimized experimental paradigms are thus required.

A third advantage of technical nature is that M/EEG and fMRI data need not be acquired simultaneously (Eichele et al. 2005; Debener et al. 2006; Huster et al. 2012) but can be acquired consecutively (for the case of EEG-fMRI). This avoids mutual interference during acquisition, and allows imaging modality-specific experimental design optimization. Note this is also a disadvantage: currently our approach does not use the information provided by single-trial based analysis of simultaneously acquired EEG-fMRI data. However, our entire approach extends directly to EEG without modification, and can thus be applied to simultaneously acquired EEG-fMRI data. Future research may address the combination of information from single-trial based analysis with representational similarity analysis as used here.

Finally, a specific limitation of our approach is a particular case of ambiguity: if two regions share similar representational relations but are activated at different time points, we cannot ascribe a unique activation profile to each. While this case cannot be excluded, it is highly unlikely due to the strong non-linear transformations of neural information across brain regions, and if observed, it is an interesting finding by itself worthy of further investigation.

### Summary statement

In conclusion, our results provide a spatio-temporal integrated view of neural dynamics underlying visual object recognition in the first few hundred milliseconds of vision. The MEG-fMRI fusion approach bears high sensitivity and specificity, promising to be a useful tool in the study of complex brain function.

## ACKNOWLEDGEMENTS

This work was funded by National Eye Institute grant EY020484 and a Google Research Faculty Award (to A.O.), the McGovern Institute Neurotechnology Program (to D.P. and A.O.), a Feodor Lynen Scholarship of the Humboldt Foundation (to R.M.C), and was conducted at the Athinoula A. Martinos Imaging Center at the McGovern Institute for Brain Research, Massachusetts Institute of Technology. We thank Chen Yi and Carsten Allefeld for helpful comments on the research.

